# Engineering of Chimeric Antigen Receptor T Cells with integrin αEβ7 Results in Augmented Therapeutic Efficacy against E-cadherin positive tumor

**DOI:** 10.1101/727446

**Authors:** Hongxing Sun, Shan He, Lijun Meng, Ying Wang, Hanghang Zhang, Yongnian Liu, Jian Wang, Min Tao, Stefan K. Barta, Essel Dulaimi, Henry Fung, Jean-Pierre J. Issa, Lei-Zhen Zheng, Yi Zhang

**Author notes:** Correspondence: Yi Zhang, MD, PhD, Fels Institute for Cancer Research and Molecular Biology, Department of Microbiology and Immunology, The Lewis Katz School of Medicine, Temple University, Philadelphia, PA 19140, USA.

## Abstract

Integrin αEβ7 (CD103) can interact with E-cadherin and promote T cell retention in epithelial tissue. However, whether the expression of CD103 on chimeric antigen receptor (CAR)-T cells may augment T cell anti-tumor activity remains unknown. Using a preclinical model, we demonstrate that CD103 engineering of human CAR-T cells significantly improves their therapeutic effects on eliminating pre-established E-cadherin expressing tumor cells in immune deficient NOD.*scid.Il2Rγcnull* (NSG) mice. Human T cells that were engineered with CAR containing 4-1BB and CD3zeta intracellular signaling domains (named BBz) expressed reduced level of CD103 in mice model. Ex vivo assays confirmed the effect of 4-1BB on repressing CD103 expression in CAR-T cells. On the other hand, we generated CD103 expressing CAR-T cells by introducing the αE gene into the CAR structure (named CD103-BBz CAR-T cells). As compared to BBz CAR-T cells, CD103-BBz CAR-T cells produced higher levels of IL-2 and underwent greater expansion in cultures and acquired greater capacity to control the growth and metastasis of E-cadherin expressing lymphoma cells in NSG mice. This effect of CD103-BBz CAR-T cells was associated with their increased capacity to infiltrate into the tumor and persist in vivo, leading to significantly improved overall survival of lymphoma mice. Our findings suggest that engineering tumor-reactive T cells with CD103 may represent a novel strategy to improve adoptive T cells anti-tumor efficacy, and this strategy may have broad implication in the epithelial solid tumor treatment.

**Highlights:**

- CAR-T cells with 4-1BB costimulatory domain express reduced level of CD103
- 4-1BB signaling antagonist TGF-β1 induced CD103 expression
- Ectopically expression of CD103 on CAR-T cells enhanced their anti-E-cadherin positive tumor capacity

Graphical Abstract:
**Graphic Summary:** The co-stimulatory molecule 4-1BB within the CAR protein potently suppresses CD103 expression. Engineering CAR-T cells with CD103 significantly enhances their capacity to proliferate and infiltrate into the solid tumor, leading to augmented anti-tumor immunity.

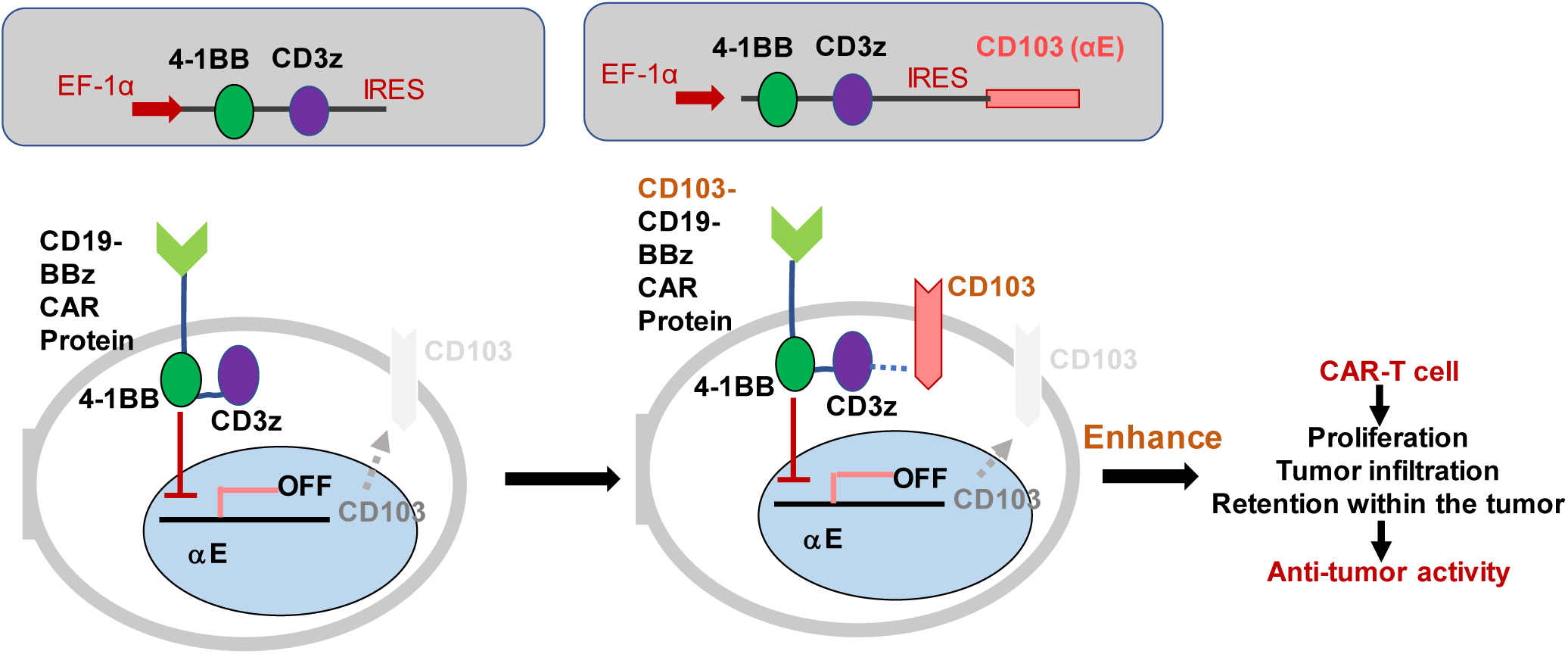

## INTRODUCTION

Whether tumor-reactive T cells can infiltrate into the tumor to execute effector function is essential for controlling tumor growth. CD103 (also known as integrin α_E_β_7_) is an integrin protein which mainly expressed on the surface of intraepithelial lymphocyte (IEL) T cells in the gut and other epithelial compartments such as skin or lung (Cerf-Bensussan et al., 1987; Kilshaw and Murant, 1991). CD103 can interact with epithelial maker-E-cadherin and promote T cell retention in epithelial tissues. Previous studies have demonstrated that CD103 is a promising marker for rapid assessment of tumor-reactive T cells infiltrating in the tumor from cancer patients, such as cervical cancers (Komdeur et al., 2017), bladder carcinoma (Wang et al., 2015), lung cancer (Djenidi et al., 2015), adenocarcinoma (Workel et al., 2016), and ovarian cancer (Webb et al., 2014). CD103 is thought to be important for retention and formation of **<u>r</u>**esident **<u>m</u>**emory T cells (T_RM_) in the peripheral tissues (Cepek et al., 1994; Mackay et al., 2013). These CD103-positive T_RM_ are found to be more prone than those CD103-negative T cells to immune checkpoint inhibition therapy (Ganesan et al., 2017; Savas et al., 2018). However, whether CD103 expression on the surface of tumor-reactive T cells is functionally crucial for eliminating tumor in vivo has not been previously determined.

Adoptive immunotherapy by engineering human T cells with a chimeric antigen receptor (CAR) specific for the pan-B-cell CD19 antigen (CD19 CAR-T cells) has been associated with high response rates in patients with relapsed or refractory B-cell malignancies, including acute lymphoblastic leukemia (ALL), chronic lymphocytic leukemia (CLL) and B-cell non-Hodgkin’s lymphoma (NHL) (Grupp et al., 2013; Kochenderfer et al., 2015; Kochenderfer et al., 2017a; Kochenderfer et al., 2017b; Locke et al., 2017; Maude et al., 2014; Porter et al., 2011a; Schuster et al., 2017; Turtle et al., 2016). Transfer of CD19 CAR-T cells induces long-lasting, complete remission in children with B-ALL and in adult patients with B-CLL. The remission is associated with high levels of in vivo expansion and persistence of CD19-specific CAR-T cells (Grupp et al., 2013; Kalos et al., 2011). However, unlike the hematologic malignancies, the efficacy of solid tumor treatment using CAR T cells is still far more satisfied (Newick et al., 2017). Since over 80% solid tumors derived from epithelial tissues and express E-cadherin, we hypothesized that engineering CAR-T cells with CD103 may augment their anti-E-cadherin positive tumor activity.

We herein demonstrate that the costimulatory molecule 4-1BB represses the induction of CD103 in CAR-T cells. Ectopic expression of CD103 in human CAR-T cells significantly enhances their anti-E-cadherin positive tumor activity. Our findings suggest that engineering tumor-reactive T cells with CD103 may represent a novel strategy to improve their anti-tumor efficacy.

## MATERIALS AND METHODS

### Mice

Immune deficient (NOD.*scid.Il2Rγcnull*) NSG mice, which lack T cells, B cells and NK cells, were purchased from The Jackson Laboratories and maintained at the Animal Facility, Temple University. Experimental protocols were approved by the Temple University’s Committee on Use and Care of Animals.

### Cell lines

The human Cell line K562 and Raji cells were maintained in RPMI 1640 supplemented with 10% FCS and 100U/ml penicillin and 100 μg/ml streptomycin sulfate. CD19 expressing K562, E-cadherin or firefly luciferase expressing Raji cells were generated by infecting them with lentiviral vector encoding CD19, E-cadherin and firefly luciferase, respectively, followed by purification using BD FACSAriaII cell sorter.

### Antibodies

Antibodies for flow cytometric analysis were purchased from Biolegend, BD Bioscience and eBioscience, including CD4-BV605, CD8-Pacific blue, CD27-BV450, CD62L-APC, CD45RO-APC-Cy7, CD45RA-PE-Cy7, CD25-PE-Cy7, CD103-APC, CD45-APC/PE/PE-Cy7, CD3-PE/APC, TCR-APC, IFN-Y-PE/PE-Cy7, IL-2-APC, CD19-PE/APC, Streptavidin-PE/APC, Eomes-PE-Cy7, T-bet-PE.CD107a-PE. E-cadherin-Biotin was purchased from eBioscience. Western blot antibodies including anti-p-Smad2, anti-Smad2, anti-β-actin were purchased from Cells Signaling Technology.

### Lentivirus Generation

pLenti-CMV-GFP (pLU) vector was acquired from Addgene (Cambridge, MA). CMV promoter was replaced by EF1a promoter to improve the expression of the encoded gene in human T cells. Lentivirus was generated by calcium phosphate transient transfection of 293T cells using psPAX2 packaging (Addgene, Cambridge, MA, USA) and pMD2.G (Addgene) envelope plasmids as described (Li et al., 2012). Lentiviral vector was concentrated by ultracentrifugation on a 20% sucrose cushion for 3 hours at 26000 rpm, 4°C.

### Isolation, transduction of primary human T cells

Whole blood was obtained from normal healthy donors, which was approved by the Institutional Review Board of Temple University. Peripheral blood mononuclear cells were isolated by Ficoll (GE Life Sciences, Marlborough, MA, USA) gradient. CD4 and CD8 T cells were isolated using positive magnetic bead-based purification (Miltenyi). Purified human T cells were seeded in 24 wells at a density of 1 × 10^6^/ml in RPMI1640 medium containing 10% FBS, 1% penicillin-streptomycin and 100U/ml IL-2. Cells were activated with Human T-Activator CD3/CD28 Dynabeads (Thermo Fisher, Waltham, MA, USA). T cells were transduced with lentiviral vector by centrifugation at 3000 rpm for 90 mins at 30°C at day 2 and day 3 after activation. Transduction efficiency was assessed at day 7 of culture. For CAR T cell purification, T cells were incubated with biotin conjugated goat anti-mouse IgG, F(ab’)_2_ fragment specific antibody (Jackson Immuno Research Labs) for 30 min at 4°C, washed, incubated with streptavidin-conjugated microbeads (Miltenyi) for additional 30 min at 4°C. Cells were separated using MACS LS column.

### Western blot analysis

Cell lysates were prepared in RIPA buffer (Thermo Fisher scientific) in the concentration of 80ul lysis buffer per 1 × 10^6^ cells. Equal amount of cell lysates (10 μl) were loaded in 4–12% Mini-PrROTEAN TGX Precast protein Gels (Bio-rad) and transferred to PVDF membrane. Membranes were blocked in 5% skimmed milk for 1 hour, incubated with the primary antibody overnight at 4 °C, washed, and incubated with the appropriate horseradish-conjugated secondary antibody for 1 hour at room temperature. An enhanced chemiluminescent (ECL) kit was used to visualize the signal (Thermo Fisher).

### Flow cytometry analysis and cytokine production

CAR-T cells were detected using biotin conjugated goat anti-mouse IgG, F(ab’)_2_ fragment specific antibody (Jackson Immuno Research Labs, PA) followed by staining with streptavidin (PE or APC). Samples were analyzed on LSRII. To induce antigen-specific production of cytokines, we co-cultured CAR-T cells with live Raji cells at 1:1 ratio for 1 hour, followed by addition of Golgi Stop (BD Biosciences) and cultured for additional 4 hours. Intracellular cytokine staining was performed to examine the production of cytokines in these CAR-T cells using Fixation/Permeabilization Solution Kit (BD Biosciences). CD107 expression was determined as described (Liu et al., 2015).

### Tumor infiltrating T cell analysis

Tumor infiltrating T cells were isolated as previously with slight modification (Watkins et al., 2012). Briefly, freshly dissected tumor masses were sheared into small pieces and digested in PBS supplemented with 10% FBS, collagenase type I (400 μg/ml) and DNase (10μg/ml) at 37 °C with stirring. Cell suspension was vortexed and filtered through 70 mm cell strainer. Cells were then purified with Percoll gradient (40% and 80%) centrifugation, washed twice with PBS supplemented with 2% FCS and analyzed using flow cytometer.

### CAR T cell cytotoxicity assay

CAR-T cells were cocultured with K562 cells containing 70% CD19^+^GFP^+^ K562 cells (on-target) and 30% CD19^−^GFP^−^ K562 cells (off-target) in indicated ratio. Mixed cultures were incubated for 48 hours and analyzed by flow cytometry. Target K562 cells co-cultured with non-CAR-T cells were used as controls.

### Induction and treatment of human leukemia and lymphoma in NSG mice

For intravenous inoculation model, 6-10 weeks old NSG mice were injected intravenously with 1×10^6^ Raji cells at day 0. CAR-T cells (0.5×10^6^ CD4 T cells + 0.5×10^6^ CD8 T cells) were given via the tail vein at day 4 post tumor inoculation. The tumor burden was monitored with PerkinElmer’s IVIS® imaging system. For lymphoma model, 1×10^6^ Raji cells or Raji cells expressing E-Cadherin (Raji-Ecad) were injected subcutaneously in the lower left quadrant abdomen. When tumor size reach 400-500 mm^3^ (about 21-25 days), 1.0 ∼ 1.5 ×10^6^ CD4 CAR-T cells were administrated intravenously. Tumor size were determined every other day. Mice developed hind-limb paralysis or tumor reach 5000 mm^3^ were terminated as endpoint.

### Statistical analysis

All results were expressed as means ± SEM as indicated. P value was calculated using Student’s *t*-test. Survival of tumor-bearing mice was analyzed using log-rank (Mantel–Cox) test. P values <0.05 were considered significant.

## Results

### BBz CAR-T cells expressing reduced level of CD103

To determine the role of CD103 in CAR T cell-mediated elimination of tumor in vivo, we first examined CD103 expression in CD19-specific CAR with 4-1BB and CD3zeta intracellular signaling domains (named CD19-BBz CAR). Using codon optimization method, we synthesized CD19-BBz CAR and cloned it into lentiviral vector, infected human T cells and produced human CD19-BBz CAR-T cells (Figure 1A, B), as previously described (Imai et al., 2004; Porter et al., 2011a; Porter et al., 2011b). *In vitro* assay confirmed the cytolytic activity of CD19-BBz CAR-T cells against K562 cells expressing of CD19 antigen (Figure 1C). To test the effect of these CD19-BBz CAR-T cells on eliminating human B-leukemic cells in vivo, we injected human lymphoma cell line Raji cells into NSG mice. Three days later when leukemia was established (Figure 1D), we transferred CD19-BBz CAR-T cells, including 0.5×10^6^ CD4 T cells and 0.5×10^6^ CD8 T cells, into these mice. As expected, CD19-BBz CAR-T cells inhibited the leukemia growth and protected these leukemic mice, with about 60% of them surviving over 60 days (Figure 1E). These results confirm the potent effect of human T cells engineered with our codon-optimized CD19-BBz CAR on eliminating B cell leukemia. Since xeno-graft versus host disease (GVHD) developed in these mice surviving the CAR-T cell therapy 50 days after infusion of CAR-T cells (data not show), we terminated these mice by day 60.

**Figure 1.**
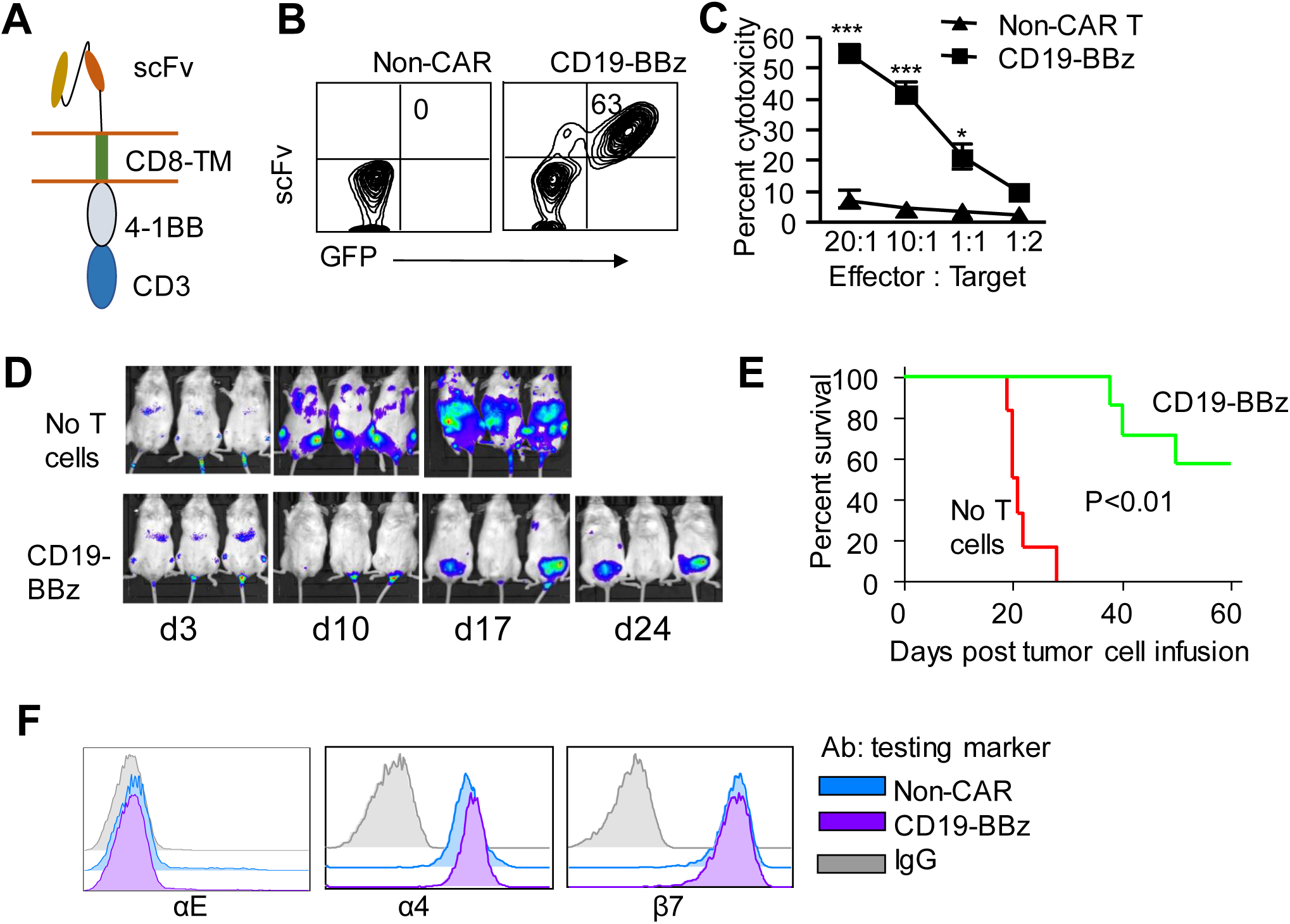
CD19-BBz CAR-T cells effectively eliminate tumor cells in NSG mice. (**A**) BBz19 CAR structure used in this study. (**B**) Expression of CD19-BBz CAR in human T cells. (C) In vitro cytotoxicity assay of CD19-BBz CAR-T cells. CD19^+^ K562 and CD19^−^ K562 cells were mixed at a ratio of 7:3, cocultured with CAR-T cells and non-CAR-T cells at indicated effector to target ratios. Twenty-four hours later, cells were collected to measure the lysis activity of CAR-T cells. Data are presented as mean ± SEM. Results are representative of three independent experiments. (**D, E**) Raji cells were injected intravenously. Three days later, 1∼1.5 ×10^6^ CD19-BBz CAR-T cells were injected via tail vein. Tumor burden were monitored by IVIS spectrum (**D**) and survival was monitored over time (**E**). (**F**) flow cytometric analysis of αE, α4 and β7 expression on the surface of T cells. *, p<0.05, ***, p<0.001.

We next measured CD103 expression on the surface of CAR-T cells. Both activated non-CAR-T cells and CD19-BBz CAR-T cells expressed similar levels of α4 and β7 integrin but no αE (Figure 1F). Upon adoptive transfer into NSG mice with pre-established Raji leukemia, approximately 5% to 10% of non-CAR-T cells produced substantially high levels of CD103 (Figure 2A). In contrast, CD19-BBz CAR-T cells failed to upregulate CD103 in the bone marrow, spleen and liver (Figure 2A). These data indicate that BBz CAR may influence the upregulation of CD103 on the surface of human T cells in vivo in these NSG mice.

**Figure 2.**
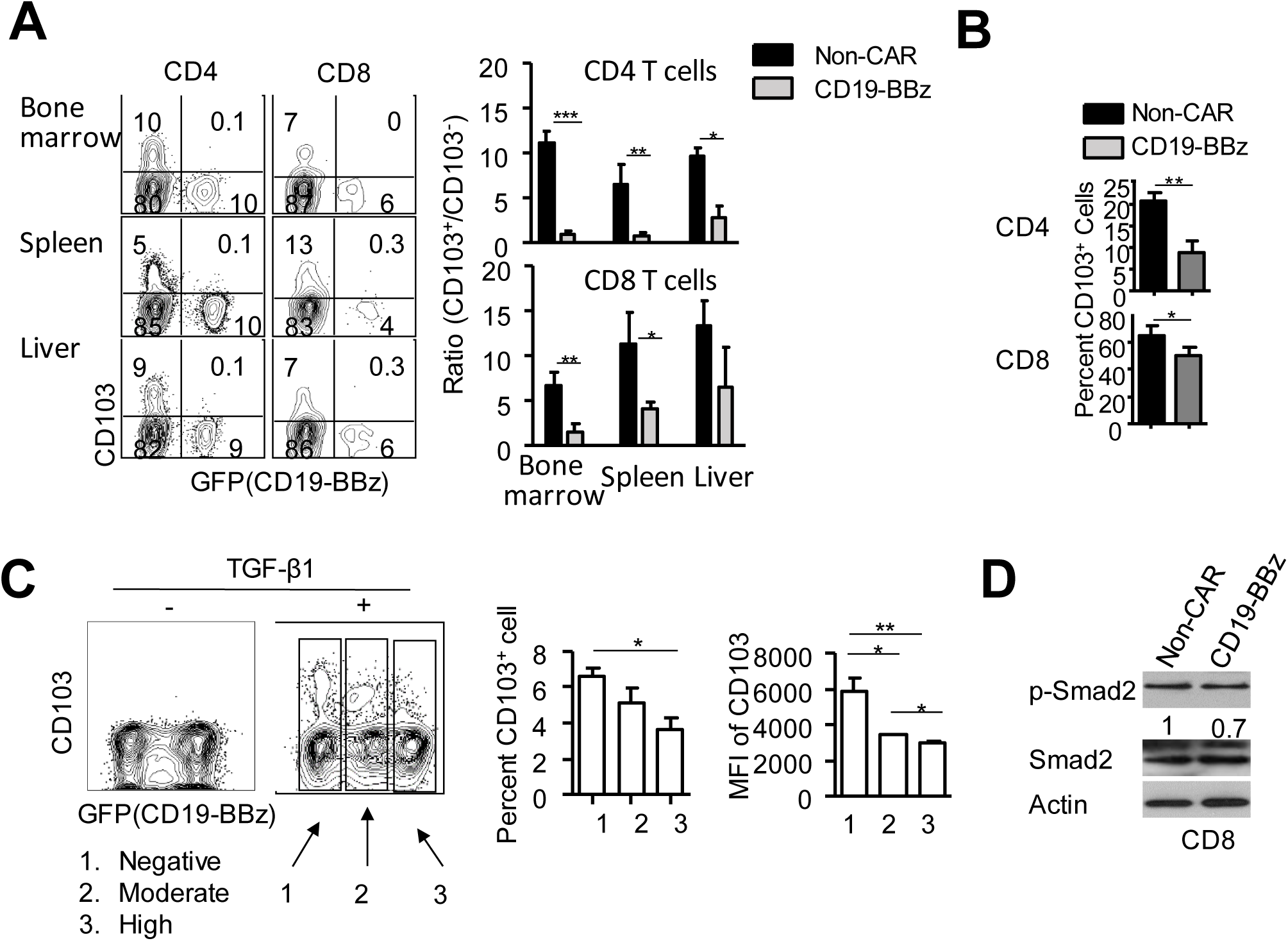
BBz CAR-T cells are unable to upregulate CD103. (**A**) Raji cells were injected intravenously on day 1. Seven days later, 1∼1.5 ×10^6^ CD19-BBz CAR-T cells were given intravenously. Mice (5/group) were euthanized on day 25 and T cells derived from different organs were collected to measure the expression of CD103. (**B**) non-CAR and CD19-BBz CAR-T cells were cultured in wells of a 24 well plate, stimulated with anti-CD3Ab for 24 hours, with or without addition of TGF-β1 (10ng/ml). Graphs show the frequency of CD103. (**C**) CD4 CAR-T cells were treated with TGF-β1 (10ng/ml) for 10 days, and the GFP and CD103 expression were determined by flow cytometry. (**D**) Smad2 expression and phosphorylation assay. Purified CAR-T cells were cultured in the present of TGF-β1 (10ng/ml) for 48 hours, followed by Western Blot. Data are presented as mean ± SEM of three donors. *, p<0.05, ***, p<0.001.

### 4-1BB signaling suppresses TGF-β1-induction of CD103 expression

Previous studies have demonstrated that TGF-β1 is important for inducing CD103 in T cells (Mokrani et al., 2014). Indeed, we found that while human T cells cultured in the presence of anti-CD3Ab/CD28 Ab-conjugated beads did not express CD103, addition of TGF-β1 readily induced CD103 on the surface of those T cells without expressing CD19-BBz CAR (Figure 2B). Notably, the down-regulation of CD103 was in parallel to the increase of the level of CD19-BBz CAR in these cultured T cells (Figure 2C). Previous studies have shown that phosphorylation of SMAD2 (p-SMAD2) is the direct downstream event of TGF-β signaling (Massagué, 2000). We found that activated CD8 CD19-BBz CAR-T cells had lower levels of p-SMAD2 compared to activated CD8 T cells without expressing the CAR structure (Figure 2D). These results suggest that CD19-BBz CAR signaling suppresses the induction of cell surface integrin αEβ7 in human T cells.

To understand the underlying mechanism by which CD19-BBz CAR-T cells expressed reduced level of CD103, we examined the selective contribution of 4-1BB and CD3z signaling to suppression of CD103. We generated human T cells expressing CD19 Ab scFv fused with CD3z (named CD3z CAR), in which CD3z CAR T cells lacked 4-1BB signaling. This allowed us to examine the differential contribution of TCR signaling and 4-1BB to the modulation of CD103 in CAR-T cells. We found that stimulation with CD19 antigen derived from Raji cells, which activates CAR signaling in T cells, failed to induce CD103 on the surface of non-CAR-T cells, CD3z CAR-T cells or CD19-BBz CAR-T cells (Figure 3A). Addition of TGF-β readily induced CD103 in these cells, with CD19-BBz CAR-T cells to significantly less extent (Figure 3A, B). This effect of CD19-BBz CAR on reducing CD103 expression was observed in both CD8 and CD4 T cells (Figure 3A, B). Taken all together, these findings indicate that 4-1BB signaling represses the upregulation of CD103 in CAR-T cells.

**Figure 3.**
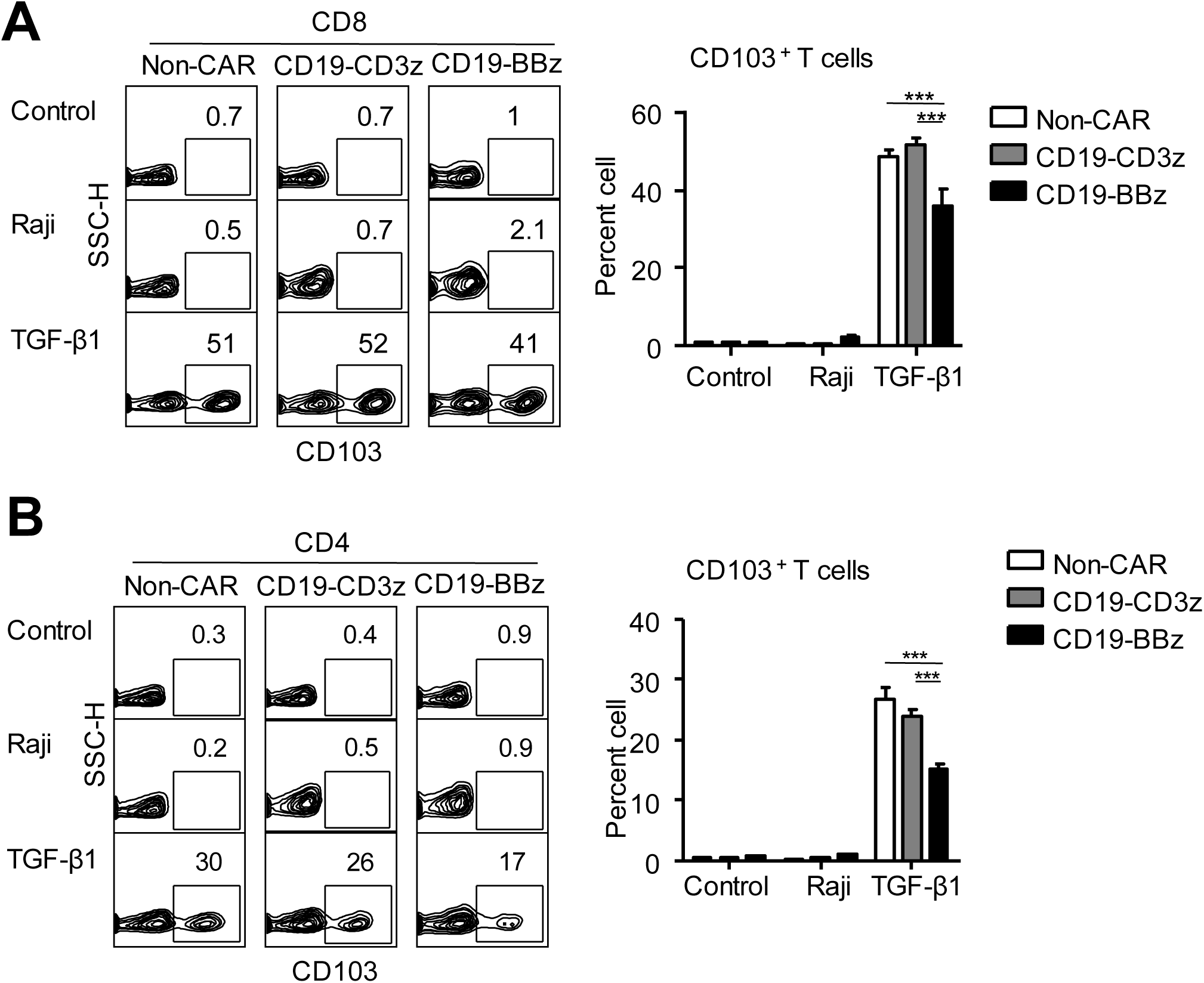
4-1BB signal suppresses TGF-β1-induced CD103 expression. (**A, B**) Normal T cells, CD19-CD3z and CD19-BBz CAR-T cells were cultured in RPMI1640 medium supplemented with 10% FBS, IL-2 (100 IU/ml), with or without addition of Raji cells or TGF-β1 (10 ng/ml). CD103 expression were determined after 48 hours coculture. T cells from three donors were used for this experiment. Data are presented as mean ± SEM of three donors. ***, p<0.001.

### Engineering of CD19-BBz CAR-T cells with CD103

To test whether CD103 expression in CAR-T cells may augment their anti-tumor activity, we introduced CD103 into CD19-BBz CAR-T cells. Since activated T cells constitutively expressed β7 integrin (Figure 1F), which binds αE (CD103) to form the αEβ7 complex, we used lentiviral vector to introduce the gene encoding αE into CD19-BBz CAR plasmid to produce CD103-CD19-BBz CAR lentiviral vector (Figure 4A). Human T cells infected with CD103-CD19-BBz CAR lentiviral vector produced both CD19-BBz and CD103 on their surface (Figure 4B, C). Overexpression of CD103 on the surface of CAR-T cells slightly affected their production of α4 and β7 molecules compared to non-CAR-T cells and conventional CD19-BBz CAR-T cells (Figure 4C). Interestingly, the majority of CD103-CD19-BBz CAR-T cells expressed were CD62L^hi^CD45RA^hi^ (Figure 4D). Since CD62L^hi^CD45RA^hi^ T cells are associated with less-differentiated T cells (Gattinoni et al., 2006). These results suggest that overexpression of CD103 may decrease the differentiation of human CAR-T cells in cultures.

**Figure 4.**
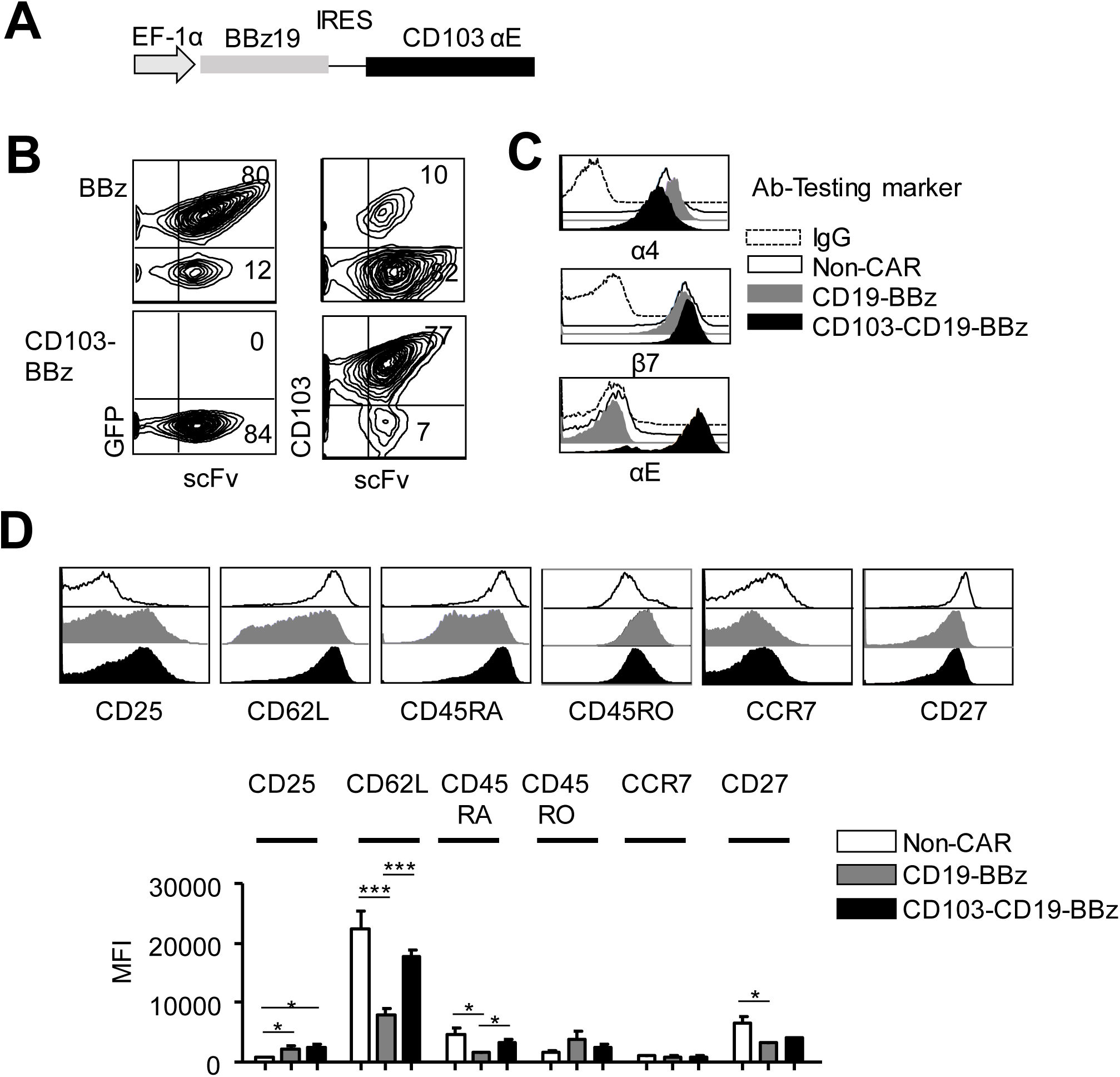
Generation and characterization of CD103-BBz CD4 CAR-T cells. (**A**) CD103 (αE) was cloned to the down-stream of IRES in the pLU plasmid encoding CD19-BBz. (**B**) CAR-T cells expressing scFv were purified on day 6 of culture, then cultured for additional 9 days and assessed for their expression of CD103 using flow cytometer. (**C, D**) Histograms show the expression of integrins and other markers on the surface of CAR-T cells. Data are presented as mean ± SEM of three donors. MFI: Mean fluorescence intensity. *, p<0.05, ***, p<0.001.

### CD103-CD19-BBz CAR-T cells have greater capacity to proliferate and produce IL-2 than conventional CAR-T cells

We then examined whether overexpression of CD103 may affect CAR-T cell proliferation and production of effector cytokines (e.g., IL-2 and IFN-γ). Both CD103-CD19-BBz CAR-T cells and conventional CD19-BBz CAR-T cells were purified at 6 days after infection, cultured for additional 15 days. As compared to conventional CD19-BBz CAR-T cells, CD103-CD19-BBz CAR-T cells underwent significant more expansion in cultures (Figure 5A). When the CAR-T cells were stimulated with Raji cells, they produced comparable CD107a (Figure 5B) but significantly higher levels of IL-2 (Figure 5C). These data indicate that overexpression of CD103 enables CAR-T cells greater capacity to proliferate and produce IL-2, which are known to be important for enhancing anti-tumor immunity in vivo (Yee et al., 2002).

**Figure 5.**
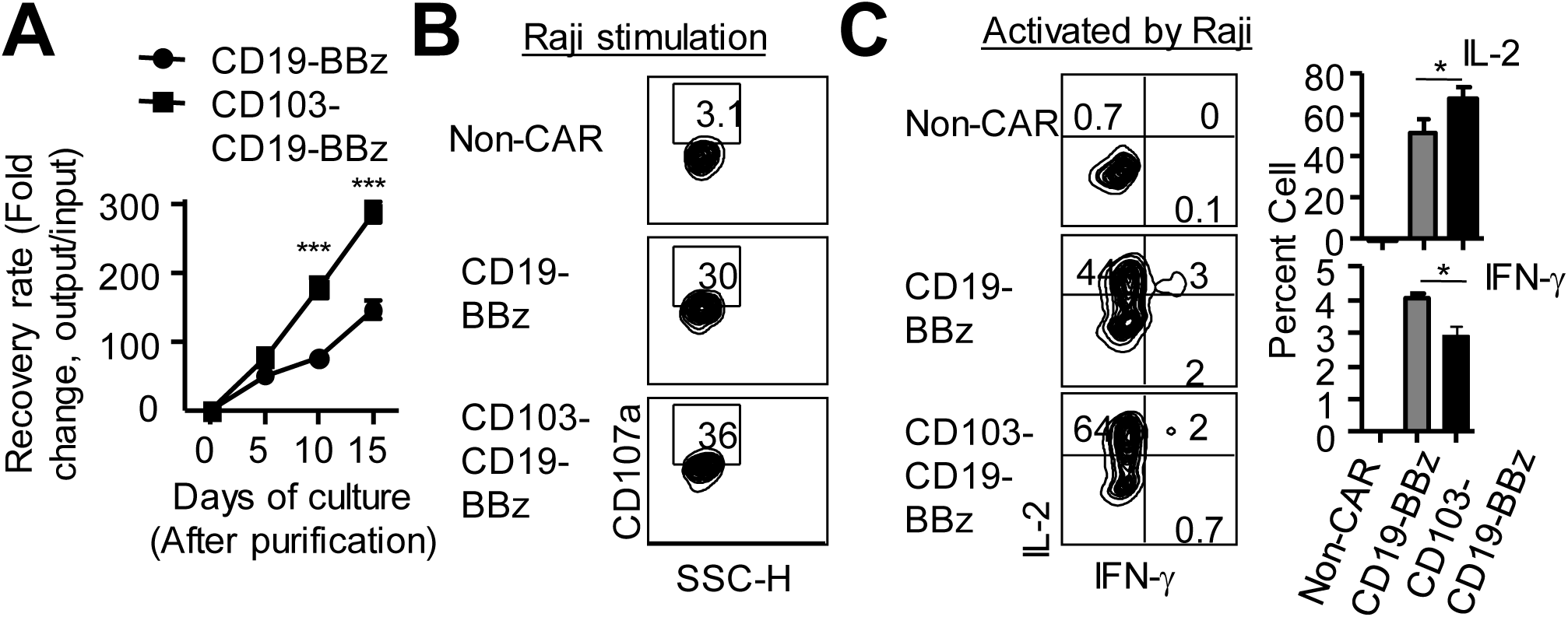
Overexpression of CD103 in CAR-T cells enhanced their proliferation and IL- 2 production. (**A**) CAR-T cells were purified on day 6 post transduction, seeded in 96 well plate and cultured for additional 15 days. Cell recovery rate was monitored over time. (**B**) Plots show the expression of CD107a in cultured T cells stimulated with Raji cells. (C) Plots and graphs show the production of IL-2 and IFN-γ by these T cells, with or without Raji cell stimulation. Data are presented as mean ± SEM of three donors. *, p<0.05, ***, p<0.001.

### The expression of E-cadherin in Raji lymphoma cells impairs the antitumor effect of conventional CD19-BBz CAR-T cells

Since most of lymphoma cells do not express E-cadherin, we thus seek to overexpress E-cadherin in Raji cell. To test whether the expression of E-cadherin may influence the susceptibility of lymphoma cells to CAR-T cell therapy, we first examined the expression of E-cadherin in several cell lines of B cell lymphoma. E-cadherin is synthesized as a 140 kDa precursor that undergoes cleavage by proprotein convertases to become a 120 kDa mature protein. This processing is essential for E-cadherin to be transported to the cell surface (Posthaus et al., 1998). Notably, both Raji and KamI cells produced E-cadherin precursor protein (Figure 6A). To induce cell surface E-cadherin, we introduced mature E-cadherin into Raji cells, named Raji-Ecad cells (Figure 6A). Subcutaneous inoculation of either Raji or Raji-Ecad cells induced lymphoma in NSG mice, with all of them dying from the disease between 35 days and 45 days after the challenge (Figure 6B). To our surprising, while adoptive transfer of conventional CD19-BBz CAR-T cells effectively controlled the growth of Raji cell-induced lymphoma, they failed to destroy the lymphoma derived from Raji-Ecad cells (Figure 6C, D). As compared to mice bearing Raji cell lymphoma, mice of Raji-Ecad lymphoma showed 2-to 3-fold less in frequency of CAR-T cells in the circulating peripheral blood at day 21 and day 28 after transfer of conventional CAR-T cells (Figure 6E), suggesting that Raji-Ecad lymphoma impaired in vivo survival and expansion of CAR-T cells.

**Figure 6.**
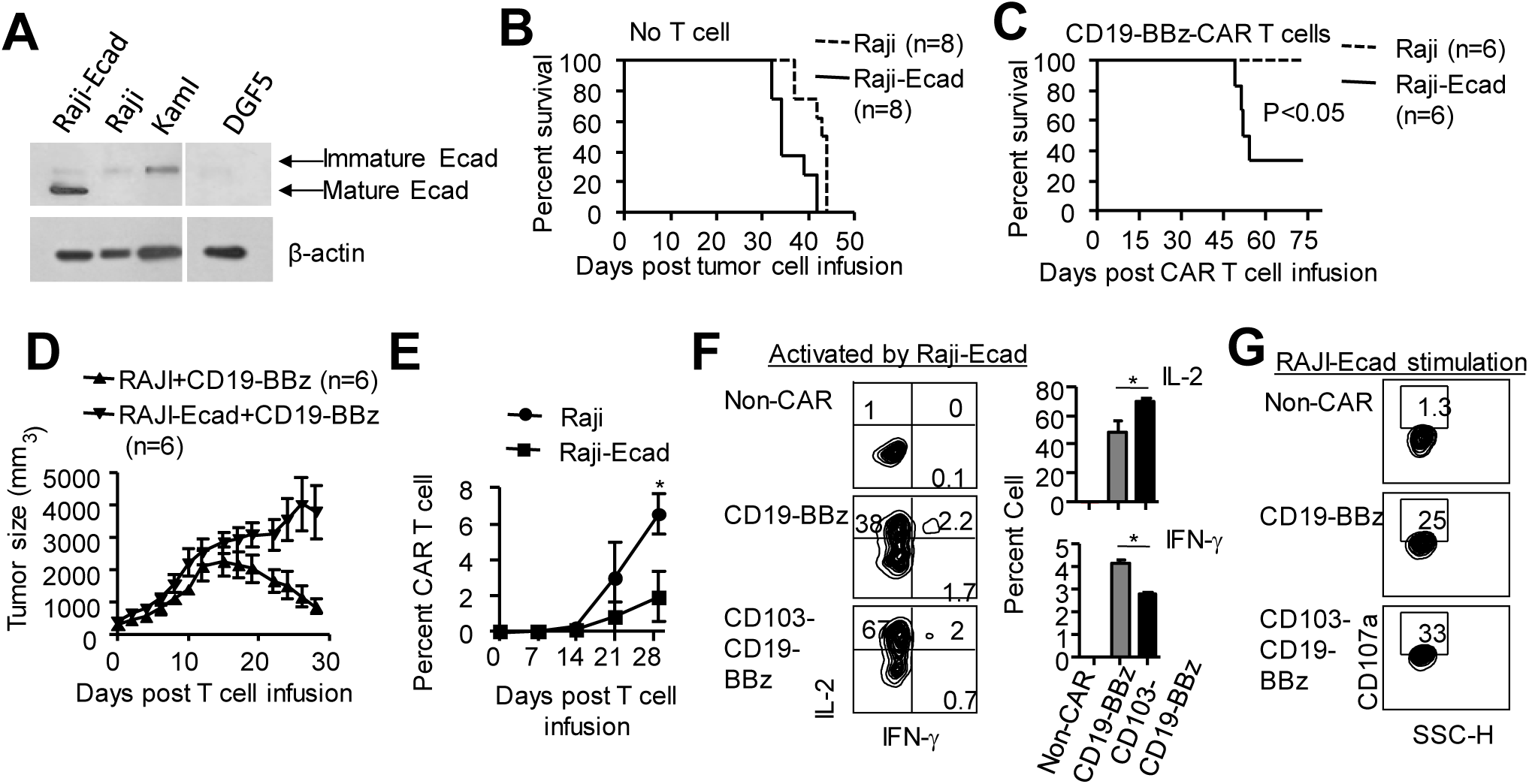
Raji-Ecad tumor cells are refractory to CD19-BBz CAR-T cell therapy. (**A**) Western blot analysis shows E-cadherin expression malignant B cells. (**B**) Equal number of Raji and Raji-Ecad cells were injected into NSG mice subcutaneously and mice survival were determined. (**C**) Equal number of Raji and Raji-Ecad cells were injected into NSG mice subcutaneously and 1∼1.5 ×10^6^ CD19-BBz CAR-T cells were administrated intravenously when tumor size reach 400-500 mm^3^. Survival (**C**) and tumor sized (**D**) were monitored. (**E**) Graph show the frequency of CAR-T cells in peripheral blood of mice challenged with Raji cells and Raji-Ecad cells, respectively. (**F, G**) CAR-T cells were cocultured with Raji-Ecad cells for 5 hours to measure cytokines (**F**) and CD107a (**G**). Plots and graph show cell percentage. Data are presented as mean ± SEM of three independent experiments. *, p<0.05.

To examine the possibility that Raji-Ecad lymphoma might impair anti-tumor activity of CD19-BBz CAR-T cells via a mechanism of decreasing their effector function, we co-incubated CD19-BBz CAR-T cells with Raji-Ecad, with or without Raji cells as control. We found that upon Raji-Ecad activation, CD19-BBz CAR-T cells significantly higher amount of IL-2- and IFN-γ-producing effectors compared to non-CAR-T cells (Figure 6F, G), but approximately 1.8-fold less IL-2-producing effector cells than CD103-CD19-BBz CAR-T cells (Figure 6F, G). These data suggest that lymphoma Raji cells expressing E-cadherin may decrease the capacity of conventional CAR-T cells to expand in vivo, leading to reduction of their anti-tumor activity. Moreover, this Raji-Ecad tumor model provided us a perfect system to test CD103-CD19-BBz CAR-T cells.

### Engineering CAR-T cells with CD103 enhances their capacity to eliminate E-cadherin positive tumor cells

Finally, we examined the impact of functional CD103 on CAR-T cell elimination of pre-established Raji-Ecad tumor in NSG mice. We transferred purified CD103-CD19-BBz CAR-T cells and conventional CD19-BBz CAR-T cells into these tumor bearing mice at day 22 to 25 after inoculation of tumor cells (Figure 7A). As compared to conventional CD19-BBz CAR-T cells, CD103-CD19-BBz CAR-T cells significantly prolonged the median survival time (69 ± 19 days versus 53 ± 11 days) and improved the overall survival rate (40% versus 10%, P<0.05. Figure 7A). This improved therapeutic efficacy of CD103-CD19-BBz CAR-T cells was associated with inhibition of tumor growth (Figure 7B), increased frequency of CAR-T cells in circulating peripheral blood (Figure 7C). Notably, CD103-CD19-BBz CAR-T cells dramatically reduced the rate of distal metastasis 80% to 30% in lymphoma NSG mice (Figure 7D). In addition, mice receiving CD103-CD19-BBz CAR T cells had approximately 2.7-fold more in frequency of CAR-T cells in the tumor than conventional CD19-BBz CAR T cells (Figure 7E). Immunohistochemistry staining indicated that the tumor infiltrating T cells showed morphological difference in terms of the cell size (Figure 7F). Altogether, the presented data suggest that the overexpression of CD103 in CAR-T cells results in significant improvement of their therapeutic efficacy in this preclinical model.

**Figure 7.**
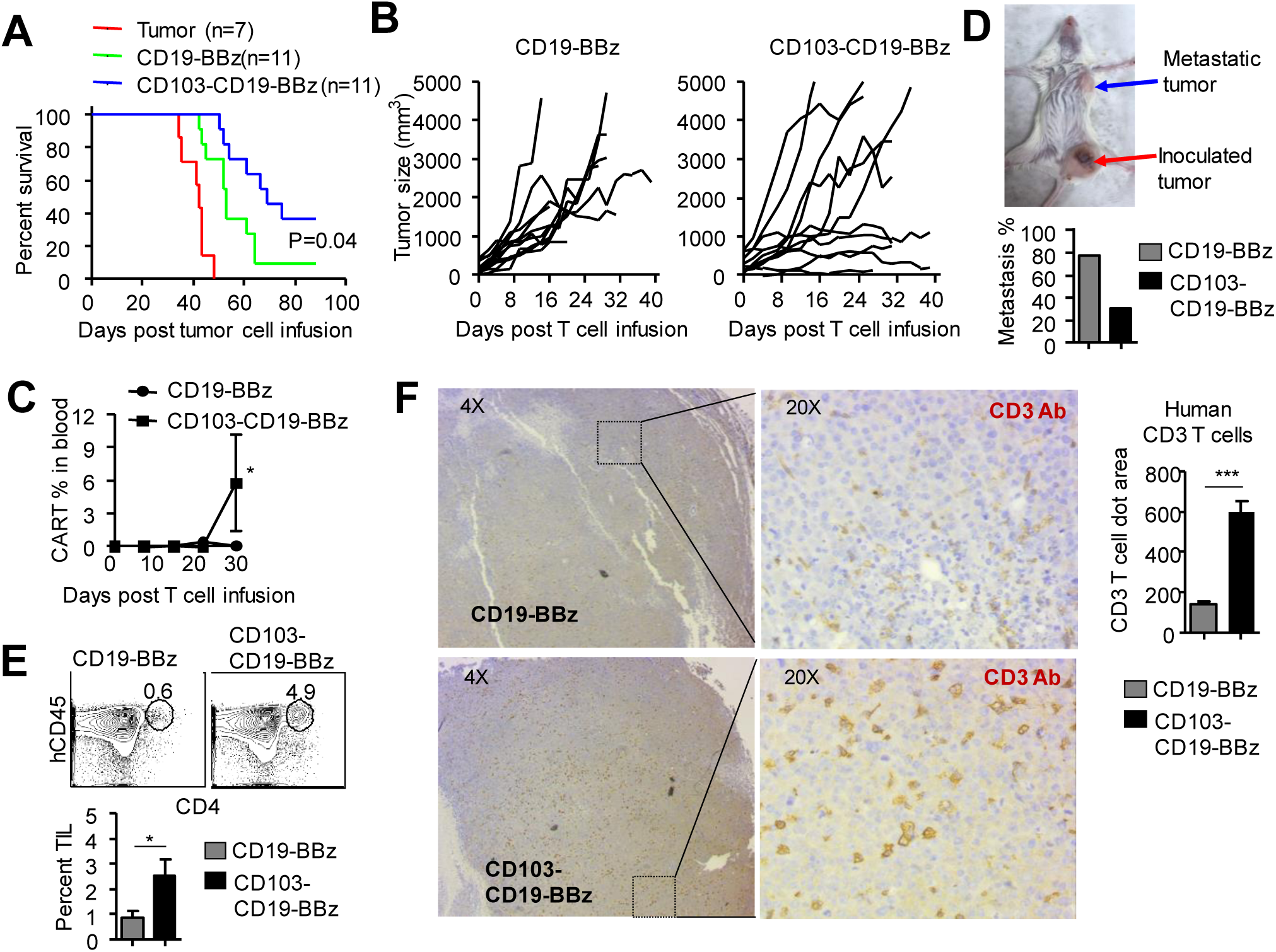
CD103-CD19-BBz CAR-T cells have greater anti-tumor activity than CD19- BB CAR-T cells. Raji-Ecad cells were injected subcutaneously at day 0 to induce lymphoma in the lower quadrant of the mouse abdomen. Twenty days later when the solid lymphoma was established, 1∼1.5 ×10^6^ CD19-BBz or CD103-CD19-BBz CD4 CAR-T cells were infused intravenously. (**A, B**) Survival of these animals and tumor size were monitored over time. (**C**) Frequency of CAR-T cells in the circulating peripheral blood. (**D**) Mice with distal metastatic tumor were monitored (n=11 for each group). (**E, F**) mice were euthanized on day 25-30 after T cells infusion. Tumor infiltrating T cells (TILs) were examined. Plots and graft show the frequency of TILs (**E**). Immunohistochemistry staining was performed to examine TILs in the tumor using anti-human CD3 antibody (**F**). Each CD3 T cell area (brown dot) were calculated with Image J and top 50 dots (based on size) were selected for analysis.

## DISCUSSION

In this study, we demonstrate that engineering CAR-T cells with CD103 augments their anti-tumor activity. These CD103-CD19-BBz CAR-T cells acquired greater capacity than conventional CAR-T cells to control the growth and metastasis of E-cadherin positive tumor in NSG mice. Overexpression of CD103 in these CAR-T cells increased their capacity to produce higher levels of IL-2 and to expand in vivo. Furthermore, CD103-CD19-BBz CAR-T cells had significantly increased capacity to infiltrate into the tumor and to persist in vivo, leading to significantly increased overall survival of mice. Our findings indicate that engineering tumor-reactive CAR-T cell with CD103 may represent a novel strategy to improve their anti-E-cadherin positive tumor efficacy and will find new application to the treatment of other types of solid tumor.

As a costimulatory factor, 4-1BB signaling induces T cell proliferation and inhibits apoptosis (Arch and Thompson, 1998; Sica and Chen, 1999). CAR-T cells expressing 4-1BB have acquired superior capability to replicate and persist in vivo (Grupp et al., 2013; Maude et al., 2014). It has been shown that 4-1BB signaling promotes replication and persistence of tumor-reactive CAR-T cells, thereby sustaining their anti-tumor activity over period of weeks and months needed to eliminate tumor (Barrett et al., 2014). However, whether persistent expression of 4-1BB may have negative impact on T cell anti-tumor activity has never been reported. The repressive effect of 4-1BB signaling on CD103 expression suggests an urgent need to better understand the mechanism by which 4-1BB signaling decreases the upregulation of CD103 in CAR-T cells. However, whether 4-1BB signaling may affect TGF-β regulation of CD103 has not been previously examined. We observed that overexpression of 4-1BB markedly decreased TGF-β-mediated upregulation of CD103 on the surface of CD19-BBz CAR-T cells. This was accompanied with inhibiting TGF-β activation of Smad2, which is known to upregulate CD103 (Mokrani et al., 2014). It will be interesting to further investigate whether selectively blocking the interaction between 4-1BB signaling and TGF-β signaling may induce CD103 expression in CAR-T cells without impairing 4-1BB effects on T cell proliferation and persistence.

One most recent study demonstrated that engineering CAR-T cells with adhesion molecule could promote T cells infiltration in brain tumor and enhance its antitumor activity (Samaha et al., 2018). CD103 is an important marker of tissue T_RM_ (Schenkel and Masopust, 2014). Earlier studies suggested that although CD103 was dispensable for the entry of T cells into the epidermis, it was crucial for T_RM_ formation(Mackay et al., 2013). CD103 was found to be important for the retention and persistence of T_RM_ in local tissues (Mackay et al., 2013). Recent studies demonstrated that tumor infiltrating T cells expressing CD103 prone to respond to immunotherapy and predicted a favorable outcome (Amsen et al., 2018; Ganesan et al., 2017; Savas et al., 2018). However, whether CD103 expression may influence the survival and persistence of antigen-experienced T cells remains largely unknown. We found that overexpression of CD103 in CD19-BBz CAR-T cells led to generation of less-differentiated CD62L^hi^CD45RA^hi^ T cells, production of high levels of IL-2 but lower levels of IFN-γ. Most importantly, overexpression of CD103 in CD19-BBz CAR-T cells dramatically enhanced their proliferation in vivo and in vitro, and their persistence within the tumor. Our findings identified a previously unrecognized function of CD103 in restraining tumor-reactive T cells from terminal differentiation and promoting the persistence of effector T cells capable of producing high levels of IL-2. This may explain, at least in part, why engineering of CAR-T cells with CD103 augments their anti-tumor activity.

E-cadherin is a calcium dependent cell adhesion molecule and mainly expressed on epithelial cells and loss of E-cadherin has been considered involved in tumor cell metastasis (Jeanes et al., 2008). The intracellular domain of E-cadherin can interact with β-catenin to repress the Wnt signaling activity, resulting in tumor growth arrest (Heuberger and Birchmeier, 2010). Thus, E-cadherin is considered a tumor suppressor. In contrast, E-cadherin expression has been shown to be associated with poor survival of glioblastoma patients. E-cadherin overexpression in glioblastoma cells increased their invasiveness in pre-clinical models (Lewis-Tuffin et al., 2010). These observations suggest that E-cadherin overexpression may influence tumor progression in a context-dependent manner. In our current study, we found that Raji lymphoma cells expressing E-cadherin significantly decreased their susceptibility to the therapeutic effect of CD19-BBz CAR-T cells in mice compared to Raji lymphoma without expressing E-cadherin. This decreased anti-tumor activity of CD19-BBz CAR-T cells in Raji-Ecad lymphoma mice was associated with markedly impaired capacity of these CAR-T cells to persist and expand in vivo. Future studies will investigate if patients with lymphoma cells expressing high levels of E-cadherin might be less susceptible to the therapy using conventional CAR-T cells, and whether CD103-CD19-BBz CAR-T cells may have augmented capacity to control tumor progression in patients with E-cadherin-positive lymphoma.

In summary, our findings indicate that ectopic expression CD103 on the surface of CAR-T cells greatly enhances their capacity to infiltration and retention within the tumor, leading to significantly improved anti-E cadherin expressing tumor capacity. Since 80% of solid tumor derived from epithelial tissues and express E-cadherin (Jeanes et al., 2008), we therefore propose that engineering CAR-T cells with CD103 may prove to be a valuable strategy to treat various types of epithelial tumors.

## Acknowledgments

This study is supported by grants of NCI (CA172106-01, Y.Z.) and NHLBI (HL127351-01A1, Y.Z.) and Fels Pilot Grant (S.H.).

## Author Contributions

H.S. and Y.Z. conceived and designed the project; H.S., S.H., Y.L., L.M, H.Z., J.P., S.B., E.AS., and Y.W. performed experiments and analyzed the data; H.S., M.T., J.W., L.Z and Y.Z designed experiments and analyzed data; H.S. and Y.Z. wrote and edited the manuscript.

